# Planar cell polarity: intracellular asymmetry and supracellular gradients of Dachsous

**DOI:** 10.1101/2022.06.18.496667

**Authors:** Adrià Chorro, Bhavna Verma, Maylin Homfeldt, Beatríz Ibáñez, Peter A. Lawrence, José Casal

## Abstract

The slope of a supracellular molecular gradient has long been thought to orient and coordinate planar cell polarity (PCP). Here we demonstrate and measure that gradient. Dachsous (Ds) is a conserved and elemental molecule of PCP; Ds forms intercellular bridges with another cadherin molecule, Fat (Ft), an interaction modulated by the Golgi protein Four- jointed (Fj). Using genetic mosaics and tagged Ds we measure Ds *in vivo* in membranes of individual cells over a whole metamere of the *Drosophila* abdomen. We find: 1. A supracellular gradient that rises from head to tail in the anterior compartment (A) and then falls in the posterior compartment (P). 2. There is more Ds in the front than the rear membranes of all cells in the A compartment, except that compartment’s most anterior and most posterior cells. There is more Ds in the rear than in the front membranes of all cells of the P compartment 3. The loss of Fj removes intracellular asymmetry anteriorly in the segment and reduces it elsewhere. Additional experiments show that Fj makes PCP more robust. Using Dachs (D) as a molecular indicator of polarity, we confirm that opposing gradients of PCP meet slightly out of register with compartment boundaries.

## Background

### (i) A brief history of PCP

“*We have, then, to imagine a system where the polarity of the cells depends on, or is, the direction of slope of a gradient”* [1].

*“It is assumed that a concentration gradient exists between the frontal and the caudal margin of the segment. In* Galleria *the scales […] orient in the direction of the steepest gradient”* [2].

Animals are largely constructed from epithelia and information about polarity within the epithelial plane is essential for organised development. For example, appendages must be built in the correct orientation, cilia must beat together in the right direction, vertebrate hairs and insect bristles must point accurately.

This process must be coordinated as fields of cells usually share the same polarity. This property is referred to as planar cell polarity or PCP [3] and the mechanisms responsible for it have been investigated by transplantation, genetics, genetic mosaics, molecular biology and modelling.

The orientation of cells must relate to the developmental landscape; where is the head? where is the midline? Does this necessary information rely on a molecular device that, with reference to embryonic anatomy, points an arrow rather as a magnetic field orients a compass needle? If so, we would need to explain how polarity information is set up in relation to the main axes of the body, how it is conveyed to the cells and how it is read locally. Long ago, Lawrence [1] and Stumpf [2] proposed independently, on the basis of experimental results in different insects, that the scalar values and slopes of morphogen gradients could provide both positional information and orienting information to the cells. A morphogen gradient was then imagined to be a supracellular concentration gradient of secreted molecules that is aligned to the axes of body or organ. The arrow of PCP would be a readout of the direction of slope of that gradient.

When research into PCP began [1–5] the genes responsible were not known but subsequently *Drosophila* genetics was applied to the problem, mutations that interfered with cell polarisation were studied and several instrumental genes identified [eg **6, 7**]. Later, genes homologous to those in *Drosophila* were found in vertebrates and elsewhere and shown to be engaged in PCP. A nice example of this conservation is the stereocilia of the vertebrate inner ear whose exact orientation is critical for balance and hearing; attempts to analyse this process have been based on studies of PCP in the fruitfly [8].

After PCP genes were identified and sequenced many have worked to understand what these proteins do. Genetic mosaics have proved to be a key method. Let’s take an early and important example: mutations in the *frizzled* (*fz*) gene cause disturbance of PCP, but what happens if a small clone of cells that lack the gene are surrounded by normal cells? Gubb and Garcia-Bellido [9] found that although the *fz^−^* clone itself produces disoriented hairs, several rows of the genetically normal cells surrounding the mutant patch were reoriented to point towards the clone, suggesting that cell interaction is a key element of the whole process.

From many years of research, it has become apparent that PCP is not directly but indirectly dependent on the slope of gradients of morphogens. Epithelial cells are oriented by gradients of other (PCP) molecules whose synthesis is regulated by and downstream of the morphogens themselves. Also, experiments have evidenced that there are two independent molecular systems of PCP; in both cell interaction is an important component and each system can act alone to polarise cells. Both systems are independently oriented by morphogen gradients [10]. These two systems may act in support or in opposition to each other. Each of these systems depend on a specific set of molecules that form bridges between adjacent cells [**10,** reviewed in **11, 12, 13**].

### (ii) The Dachsous/Fat system

Here we study one of these two molecular systems, the Ds/Ft system. Mutations affecting Ds and Ft cause misoriented cells. Their genes were found to specify large atypical protocadherin molecules. A Ft molecule in one cell is thought to bind to a Ds molecule in the other cell, thereby stabilising both molecules in the cell membranes and forming a heterodimeric bridge from cell to cell [14, 15].

Consequently, the accumulation of one molecule, say Ft, in a cell can affect the disposition of the other molecule, Ds, in the neighbouring cell — whose polarity may thus be altered, affecting the next neighbouring cell and so on [10].

The orientations of many cells are thought to be coordinated by one supracellular gradient of Ds activity. The shape of this gradient may be determined by not only the distribution of Ds protein itself but also, in the eye, by an opposing supracellular gradient of Fj [16]. Fj is a Golgi-resident kinase molecule that reduces the activity of Ds (in its binding to Ft) and increases the activity of Ft (in its binding to Ds) [17, 18]. See figure 1 for a summary.

**Figure 1.**
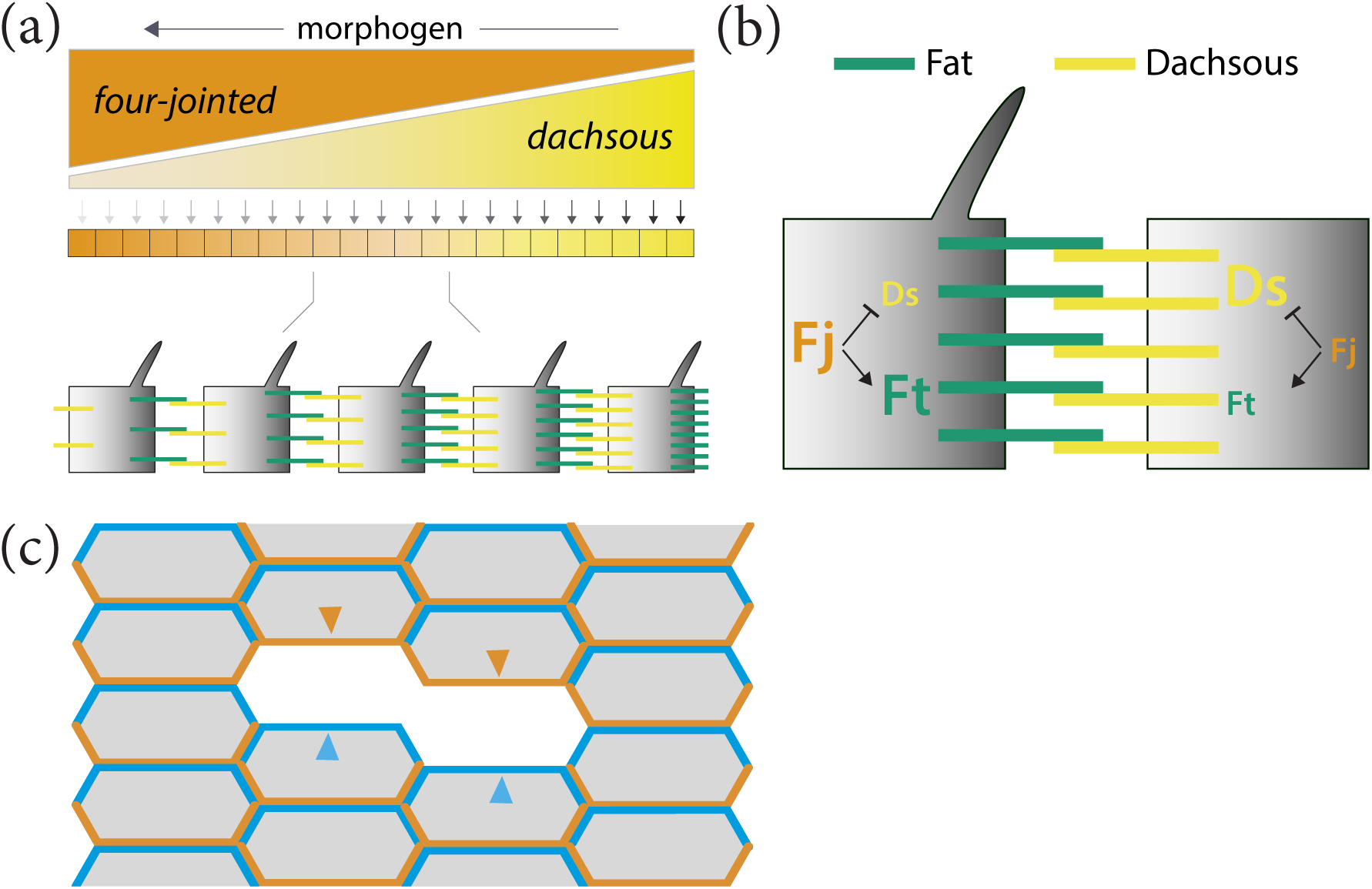
Model of the Ds/Ft system. (a, b) The anterior, or A compartment of a segment in the abdomen is shown. In response to gradient(s) of morphogen(s), opposing supracellular gradients of Fj and Ds are established. Fj predominates in the anterior region and Ds in the posterior region. Fj affects the binding of Ds with Ft and consequently both the Fj gradient and the gradient of Ds itself determine the distribution of Ds-Ft and Ft- Ds in the cells. A cell determines its polarity by comparing the disposition of Fat and/or Ds between its anterior and posterior membrane [10]. (c) How we isolate anterior or posterior membranes to measure tagged Ds in each. All the cells contain normal amounts of Ds, half of which is tagged. Tagged Ds is removed in small clones and replaced with normal untagged Ds. Consequently, the tagged Ds in either only the posterior membrane (orange) or only the anterior membrane (blue) of a cell flanking the clone can be measured.

Yang et al [19] made the first observations suggesting that, in the *Drosophila* eye, Ds and Fj are distributed in opposing gradients whose orientation relates to ommatidial polarity. Similarly, in the adult abdomen, Casal et al [20] deduced from studying mutant clones and enhancer traps that, normally, Ds is graded in opposite directions in the A and P compartments and that Fj is also graded, but in the opposite sense to Ds, in both compartments.

Evidence from enhancer traps and genetic mosaics have argued that both Ds and Fj are present in opposing gradients of function also in the eye and the wing [21, 22] but we have no precise picture of the ranges, the shapes or the steepness of the gradients in any of these organs. There is an earlier molecular investigation of Ds in the abdominal metamere which indicates that there are gradients of amount [23]. We assess this data as supportive of gradients but preliminary— their quantitation does not measure the amount of Ds on single membranes (which we believe to be necessary) but records the distribution of Ds on joint membranes. No quantified data is provided and therefore the slope of any gradients remain unknown (Fig. 4 of their paper). Also, it was assumed that the boundaries of gradients are colinear with the lineage boundaries, which as we show below was not a correct assumption.

Evidence that the Ds/Ft system can drive PCP directly — and not via the Stan/Fz system, as had been proposed [**19, 24,** reviewed in **25**] — came from experiments by Casal et al. [10] in the abdomen [reviewed in **26**]. But that finding raised the question: how does it do so? We proposed that the numbers of bridges and their orientations (Ds-Ft or Ft-Ds) differed in amounts between the anterior and posterior membranes of each polarised cell (numbers that together determine the intracellular asymmetry). That molecular asymmetry within single cells was measured by Strutt’s group but only in a small area of the wing disc near the peak of the gradient of Ds [27] and where the cells are strongly asymmetric in the localisation of Dachs. Here we assess both the intracellular asymmetry and the supracellular gradient by measuring the amount of Ds of all single membranes over a whole metamere of the abdomen. We also study and analyse the effect of Fj on these parameters and explain its role in the Ds/Ft system itself. The abdomen was chosen as it is made up of atavistic segments rather than the wing or the eye which are appendages. Finally, and this is an important advantage, only in the abdomen is there a simple relationship between the axis of PCP (the hairs and denticles point posteriorly) and the main axis of the body (the anteroposterior axis).

## Materials and Methods

### Mutations and transgenes

Flies were reared at 25°C on standard food. The FlyBase [28] entries for the mutant alleles and transgenes used in this work are the following:

*hs.FLP*: *Scer\FLP1^hs.PS^*; *tub.Gal4*: *Scer\GAL4^alphaTub84B^*; *UAS.nls-GFP*: _Avic\GFP_UAS.Tag:MYC,Tag:NLS(SV40-largeT)_; *tub.Gal80*: *Scer\GAL80*_*alphaTub84*_; *UAS.fz*:_ *fz^Scer\UAS.cSa^*; *fj^−^*: *fj^d1^*; *ds::EGFP*: *Avic\GFP^ds-EGFP^*; *CycE^−^*: *CycE^KG00239^*; *y^+^*: *Dp(1;2)sc^19^*; *w^+^*: *w^+30c^*; *en.Gal4*: *Scer\GAL4^en-e16E^*; *UAS.DsRed*: *Disc\RFP^UAS.cKa^*; hs.CD2: Rnor\Cd2^hs.PJ^; *UAS.ft*: *ft^UAS.cMa^*; *act>stop>d::EGFP*: *d^FRT.Act5C.EGFP^*.

### Experimental genotypes

***UAS.fz* clones**: *y w hs.FLP tub.Gal4 UAS.nls-GFP / y w hs.FLP; FRT42D tub.Gal80/ FRT42D pwn; UAS.fz/ +*

***UAS.fz* clones in *fj^−^***: *y w hs.FLP tub.Gal4 UAS.nls-GFP / y w hs.FLP; FRT42D tub.Gal80 fj^−^/ FRT42D pwn fj^−^; UAS.fz/ TM2*

**Untagged *ds* clones:** *y w hs.FLP; ds::EGFP CycE^−^ FRT40A / y^+^ w^+^ FRT40A en.Gal4 UAS.DsRed; +/ TM2*

**Untagged *ds* clones in *fj^−^***: *y w hs.FLP; ds::EGFP CycE^−^ FRT40A fj^−^/ y^+^ w^+^ FRT40A fj^−^ en.Gal4 UAS.DsRed*

***UAS.ft* clones:** *y w hs.FLP/ w; FRT42D pwn / FRT42D tub.Gal80 hs.CD2; UAS.ft / tub.Gal4*

***d::GFP* clones:** *y w hs.FLP/ w; +/ y^+^ w^+^ FRT40A en.Gal4 UAS.DsRed; act>stop>d::EGFP/ +*

***d::GFP* clones in *fj^−^*:** *y w hs.FLP/ w; fj^−^/ y^+^ w^+^ FRT40A fj^−^ en.Gal4 UAS.DsRed; act>stop>d::EGFP/ +*

### Cuticle clones

To induce clones overexpressing *fz* or *ft*, pupae of the appropriate genotypes were heat shocked, at 96-120 hours after egg deposition, at 37°C for 30 minutes in a water bath. 2-3 days after eclosion, adult flies were selected and kept in tubes containing 70% ethanol. Cuticles were dissected and mounted in Hoyers medium. Images were taken on a Zeiss Axiophot microscope (Carl Zeiss Ltd, Cambourne, UK) equipped with Nomarski optics using a 40x/0.90 Plan-Neofluar lens, a Nikon D-300 camera (Nikon Uk Ltd,, Surbiton, UK) connected to an iMac computer, and Nikon Camera Control Pro 2. Stacks of images taken at different focal planes were combined into a single image with Helicon Focus (HeliconSoft, Kharkiv, Ukraine).

### Quantification of polarisation strength

Overlapping images of adult cuticles containing overexpressing *fz* clones, labelled with *pawn,* were stitched together using Adobe Photoshop with the object of including the whole pigmented and haired area of the A compartment in a single image that was saved as a TIFF file. The file was opened with the ImageJ bundle Fiji. The segmented line tool was used to estimate the size of the A compartment using pigmentation and hairs as landmarks [29], to measure the average distance between the anterior boundary of the A compartment and the anterior border of the clone, and the average length of the cuticle anterior to the clone that showed reversed polarization. Due to the irregular shape of the clones the measurements were done at three different positions for each clone and the resultant average was used for the final plot. Note that for clones in the anterior [a2 region **29**] of the A compartment, measurement of effect was limited, not by the extent of repolarisation but by the lack of hairs in the *a1* region. Therefore clones close to the anterior boundary of hairs were not scored.

### Live imaging of pupal epidermis

To induce clones expressing *d::GFP* or untagged *ds* clones, pupae of the appropriate genotypes were heat shocked at 24 hours after puparium formation at 33°C for 5 or 15 minutes respectively in a water bath. 24 hours later, a 2x2 cm spacer was prepared with 7 layers of double-sided tape (Tesafix 4964, Tesa UK Ltd., Milton Keynes, UK), and a hole 6 mm in diameter was punched out of the centre; the spacer was attached to a microscope slide. Each pupa was removed from the puparium, transferred to the hole with its dorsal side facing up, covered with Voltalef 10S oil (VWR International, Lutterworth, UK) and sealed with a coverslip. Epidermal cells in the A3–A5 abdominal segments of the pupa were imaged live using a Leica SP5 inverted confocal microscope with a 63×/1.4 oil immersion objective. Tagged fluorescent proteins were excited sequentially with 488 nm and 561 nm laser beams and detected with 500 – 540 nm and 570 – 630 nm emission filters, using Leica HyD hybrid detectors. To maximise the dynamic range and avoid clipping, the pixel depth was set to 12 bits and the gain and laser power adjusted appropriately. Stacks of 1024 x1024 pixel images were thus acquired.

### Quantification of Ds

Image stacks were opened in Fiji, and projected into a single image with the Maximun intensity projection algorithm. Background was subtracted with a Rolling Ball of 6 pixels. The coordinates of the A/P and P/A boundaries (determined by the limits of *engrailed* expression) were obtained, as well of the average fluorescence intensity of the Ds signal in a 40x15 pixel box situated in a region of the A compartment free of clones and abutting the A/P boundary. Using the Freehand tool with a 6 pixel width, we measured the intensity of the Ds signal at the posterior and anterior border of untagged Ds clones (i.e. the intensity of the signal originated from a single anterior or posterior cell membrane, see Figure 1), recording the average of the intensity of three separate measures. Twin clones carrying two doses of *ds::EGFP* would be also homozygous for a *Cyclin E* mutation rendering them unable to proliferate. The coordinates of the centre of each freehand line was also obtained, allowing us to determine the position of the clone borders relative to the length of the A or P compartments. Each fluorescence intensity was standarized with respect to the intensity of the box measured before, and finally the *Relative Levels* calculated as *log(Relative Intensity) – 3*.

### Statistics and Plotting

We used RStudio with R v.4.1.2 [30], and the tidyverse, readxl, patchwork, ggpubr, tidymodels, rstatix, and mgcv packages.

## Results

There are several interrelated projects:

1. We measure the differences in Ds amounts on opposite sides of individual diploid cells (histoblasts) across the whole abdominal metamere of the living pupa. This same data tells us also how the amount of membraneous Ds varies across an entire segment, each segment comprising one A and one P compartment.
2. We study the contribution of Fj to the Ds/Ft system with respect to the metameres.
3. We map the molecular polarity of every cell in a segment using Dachs.

Here we use a normally regulated and fluorescently tagged Ds molecule [thanks to **27**]. To measure the tagged Ds in any one cell membrane its protein must be singled out from any fluorescent signal contributed by the abutting membrane of a neighbour cell — this is achieved by making many patches (clones) of cells that contain only untagged Ds, thereby isolating single membranes bearing tagged Ds that face the periphery of these clones (see Material and Methods).

### (i) Supracellular distribution of Ds across an abdominal metamere

In a single metamere of the pupal abdomen about 37 cells were counted along the anteroposterior axis from front to back of the A compartment and about 11 cells spanned the P compartment (all cell divisions having stopped by this stage).

To find the distribution of Ds, we sampled along the anteroposterior axis, these numbers are then plotted against segment length. We report a supracellular gradient in the A compartment in which Ds increases in amount towards the rear as predicted [**10, 20,** see also **23**]; a quasilinear correlation is clearly seen and is robust and statistically significant (figure 2a). We find that the Ds gradient rises steadily to achieve a difference in relative levels of 30% between its posterior and anterior limits. This quantitation confirms that the supracellular gradient of Ds in the P compartment is reversed to show a difference in amount between its anterior and posterior limits of about 15% (figure 2a).

**Figure 2.**
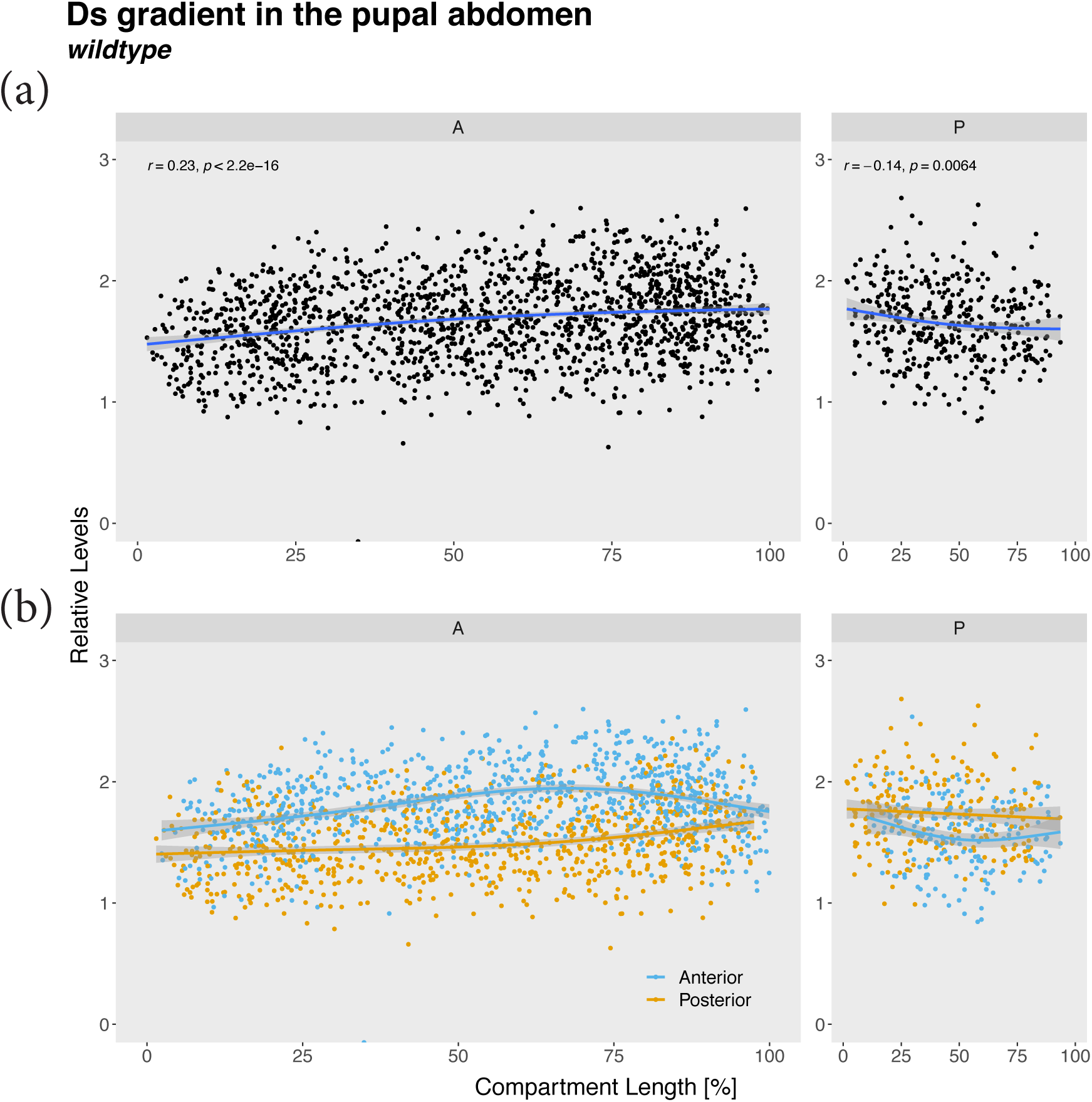
The supracellular gradient and cellular asymmetry of Ds in wildtype pupal epidermis. (a) All the measurements of Ds (both anterior and posterior cell membranes) are plotted across an entire metamere (0-100% of compartment length). A and P compartments are shown separately. The position of the compartment boundary was determined by mapping the expression of *engrailed*. Supracellular gradients are estimated to rise from the front to the back by 30% in the A compartment, falling in the P compartment by 15%. The shaded area represents the 95% confidence interval for the fitted curve. (b) The data points from above separated into anterior (blue) and posterior membranes (orange). Note both sets of data are graded but differ consistently in relative Ds amounts (but see figures S1, S2). In the A compartment the amount of Ds is greatest in the anterior membranes (peaking at about 40% near the middle of the compartment). In the P compartment the amount Ds is greatest in the posterior membranes.

### (ii) Cellular asymmetry measured across an abdominal metamere

The results are shown (figure 2b). The data for the anterior and posterior membranes are plotted separately. As predicted [10], within the A compartment the anterior membranes contain more Ds than the posterior with the relative levels changing across the segment. In the P compartment there is cellular asymmetry also but with the opposite sign (high in the posterior membrane, as expected). In the P compartment this asymmetry is statistically secure only in the central region located away from the A/P and P/A borders.

The maximum difference of relative levels of Ds between the anterior and posterior membranes occurs in the middle of each compartment being ca 40% in the A compartment and ca 20% in the P (figure 2b).

### (iii) Cellular asymmetry and supracellular gradients in the absence of Fj

Fj is clearly part of the Ds/Ft system, but the loss of Fj causes only slight effects on the wildtype phenotype. Nevertheless, comparing *fj^+^* and *fj^−^* genotypes of the A compartment we find that the cellular asymmetry is significantly reduced relative to wildtype, most clearly in the anterior 20% of the A compartment (compare figures 2b and 3b). A superposition of the data from both genotypes shows that, remarkably, the accumulation of Ds in the anterior membranes is not detectably affected by the removal of Fj (figure 2c). However, the relative levels of Ds recorded on posterior membranes of the cells is decreased in *fj^−^* as compared to the wild type (figure 2c). The same comparison in the P compartment (where, compared with the A compartment, the gradient and cellular asymmetry are reversed) shows that the loss of Fj has its largest effect also on the posterior membranes (figure 2c).

**Figure 3.**
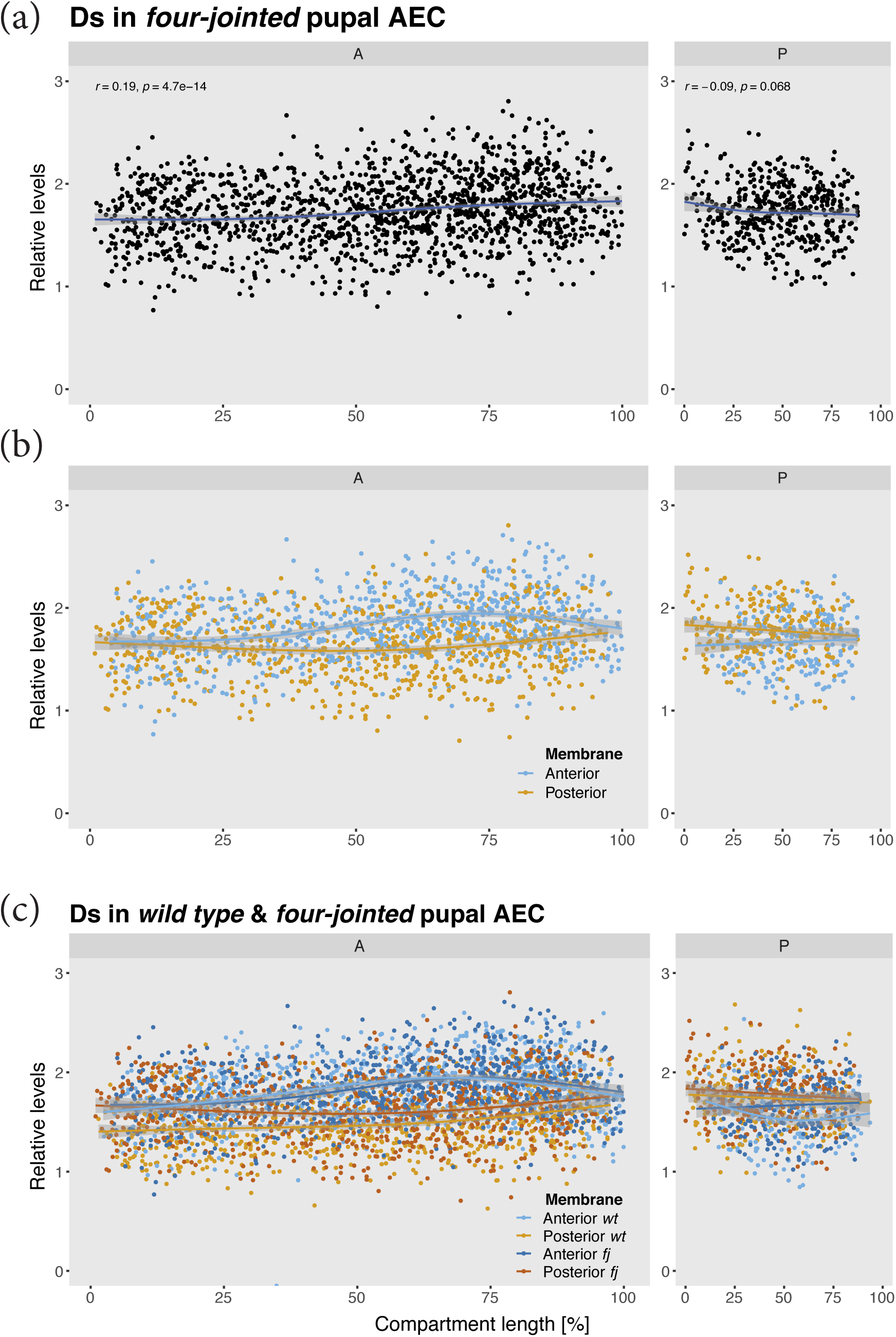
The supracellular gradient and cellular asymmetry of Ds in *fj-* pupal epidermis. (a) and (b) compare with data shown in Figure 2, the gradients are similar but somewhat shallower. In (a) note the loss of cellular asymmetry in the anterior region of the A compartment. (c) The wildtype and fj- cell asymmetry data are both shown on one plot. The anterior membranes, in both A and P compartments, have similar values in both wildtype and *fj^−^*. Note that, in the A compartment of *fj^−^* pupae, there is less Ds on the posterior membranes than in the wildtype, reducing the cellular asymmetry everywhere but especially in the anterior region.

Note that the supracellular gradients of both wildtype and *fj^−^* differ little but there appears to be some reduction in the Ds gradient in *fj^−^*, again in the most anterior region of the A compartment. (compare figures 2b and 3a).

### (iv) Estimating the effects of Fj on the robustness of the Ds/Ft system

Even though the loss of Fj mainly affects the legs, the Fj protein may still have an important function in the abdomen. It could be that Fj makes the Ds/Ft system more robust and this is evidenced by a reduction in the asymmetric distribution of Ds in the cellular membranes of *fj^−^* cells (see above). This hypothesis can be tested: we employ the Stan/Fz system to reverse polarity locally within the A compartment and to succeed it must overcome the Ds/Ft system (which is trying to maintain normal polarity). Thus, the more robust the Ds/Ft system is, the better it will be able to resist and reduce the polarising effects of the Stan/Fz system. We make small marked clones overexpressing *fz,* a key component of the the Stan/Fz system; these cause all the cells around to point their hairs away from the clones, an effect only readily apparent in those areas anterior to those clones [31]. Comparing the polarity effects of *fz*-expressing clones in various positions in the anteroposterior axis, one sees no clear trend in wildtype flies.

However, the local reversal of the polarity of bristles spreads much further in a *fj^−^* background, notably in the anterior region (figure 4a). These findings argue that that Fj strengthens the robustness of the Ds/Ft system, particularly at the front of the A compartment. This makes sense as there is indirect evidence that, in the wildtype, the amount of Fj is graded within each segment, with the highest amount anteriorly where *fj^−^* clones show the strongest phenotype [20, 21].

**Figure 4.**
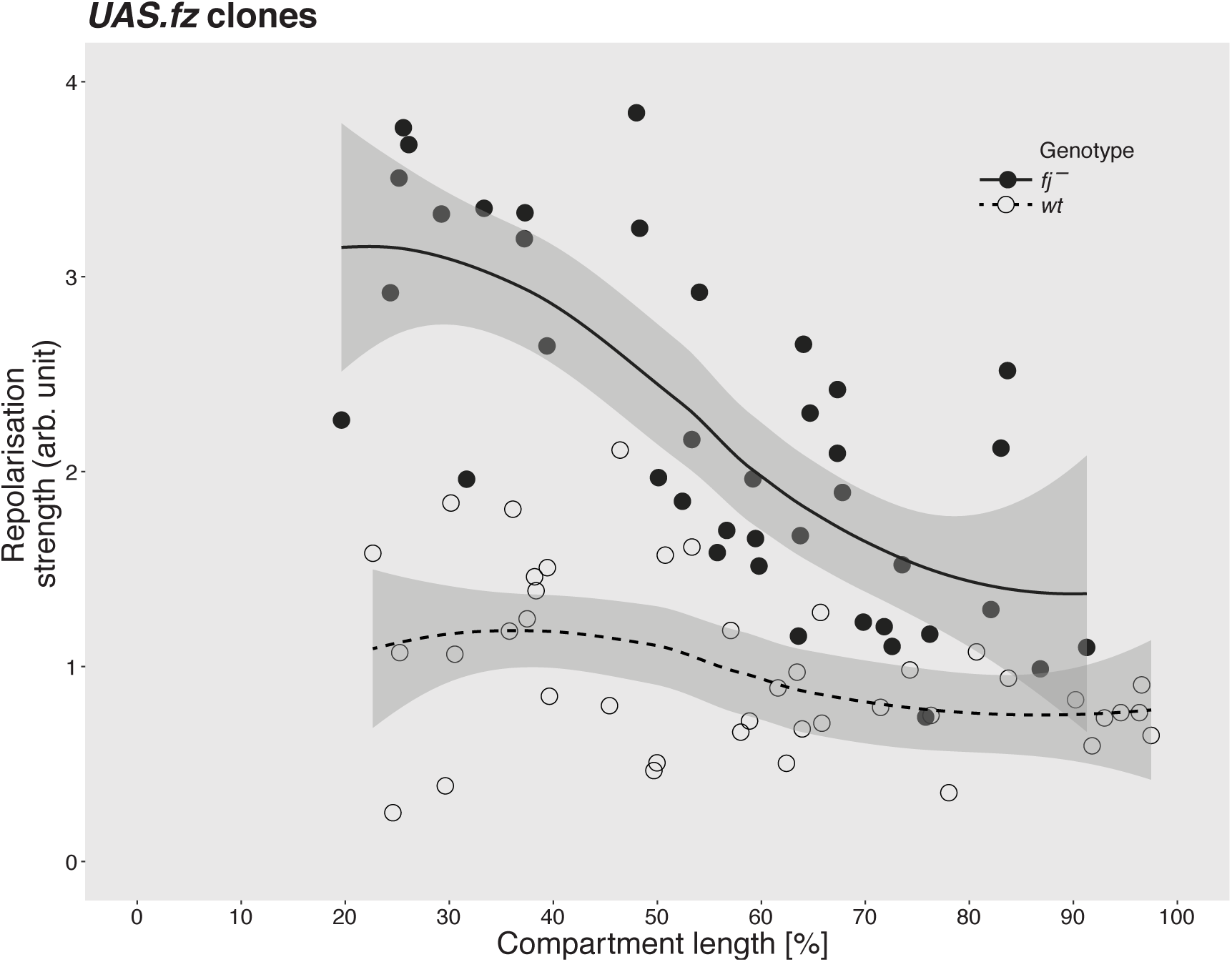
Estimating the robustness of the Ds/Ft system in the A compartment. Clones overexpressing *fz* reverse polarity anterior to the clone and they do so by overcoming the Ds/Ft system. In wildtype pupae the extent of reversal is more or less uniform within the A compartment. However, in the absence of Fj, the Ds/Ft system is weakened in the anterior regions as shown by increased range of reversal, when compared to wildtype (The shaded area represents the 95% confidence interval for the fitted curve).

### (v) Using Dachs to map cellular asymmetry throughout the pupal segment in *fj^+^* and *fj^−^* flies

The plots of Ds distribution (figure 2) showed that the cellular asymmetries dwindle and cross over near the A/P and P/A borders, not showing exactly where the two opposing gradients meet and raising uncertainty as to where cell polarity changes. Dachs (D) protein is an excellent indicator of the polarity of the Ds/Ft system [32–34] . D is located on the membrane of the cell with the most Ds and this polarity may or may not correlate with other indicators, such as the pointing of adult hairs in the P compartment [23, 35]. Given that we find most of the A compartment has a Ds gradient increasing posteriorly, are all the cells of the A compartment polarised appropriately (according to current models, D localises at the membrane where Ds is in excess [33], and therefore should be localised anteriorly in the A compartment [10]). Given that most of the P compartment has the opposite gradient, do all the cells of the P compartment have D localised posteriorly? D localisation is reported to switch from anterior to posterior polarity where the A and P compartments meet [23, 35]. However, we re-examine this by means of the distribution of D in the pupal abdomen of both wildtype and *fj^−^* flies, using small clones carrying tagged D in a background in which none of the D is tagged. In many cells it is obvious whether D accumulates mainly or only on the anterior or posterior side. The results show that in the wildtype, in **most** of the A compartment D is found in the anterior membrane (figure 5a) and in **every cell** of the P compartment D is located in the posterior membrane (figure 5b). Within the A compartment, about two rows of the most anterior cells (figure 5c) and about two rows of the most posterior cells (figure 5d) show D located posteriorly, meaning that their Ds/Ft polarity is that normally characteristic of the P cells. The larva, having fewer but larger polyploid cells told a similar story: a set of cells in the A compartment, those confined to the extreme posterior row, had variable polarity with some showing the same polarity as in the P compartment [D accumulating posteriorly **36**].

**Figure 5.**
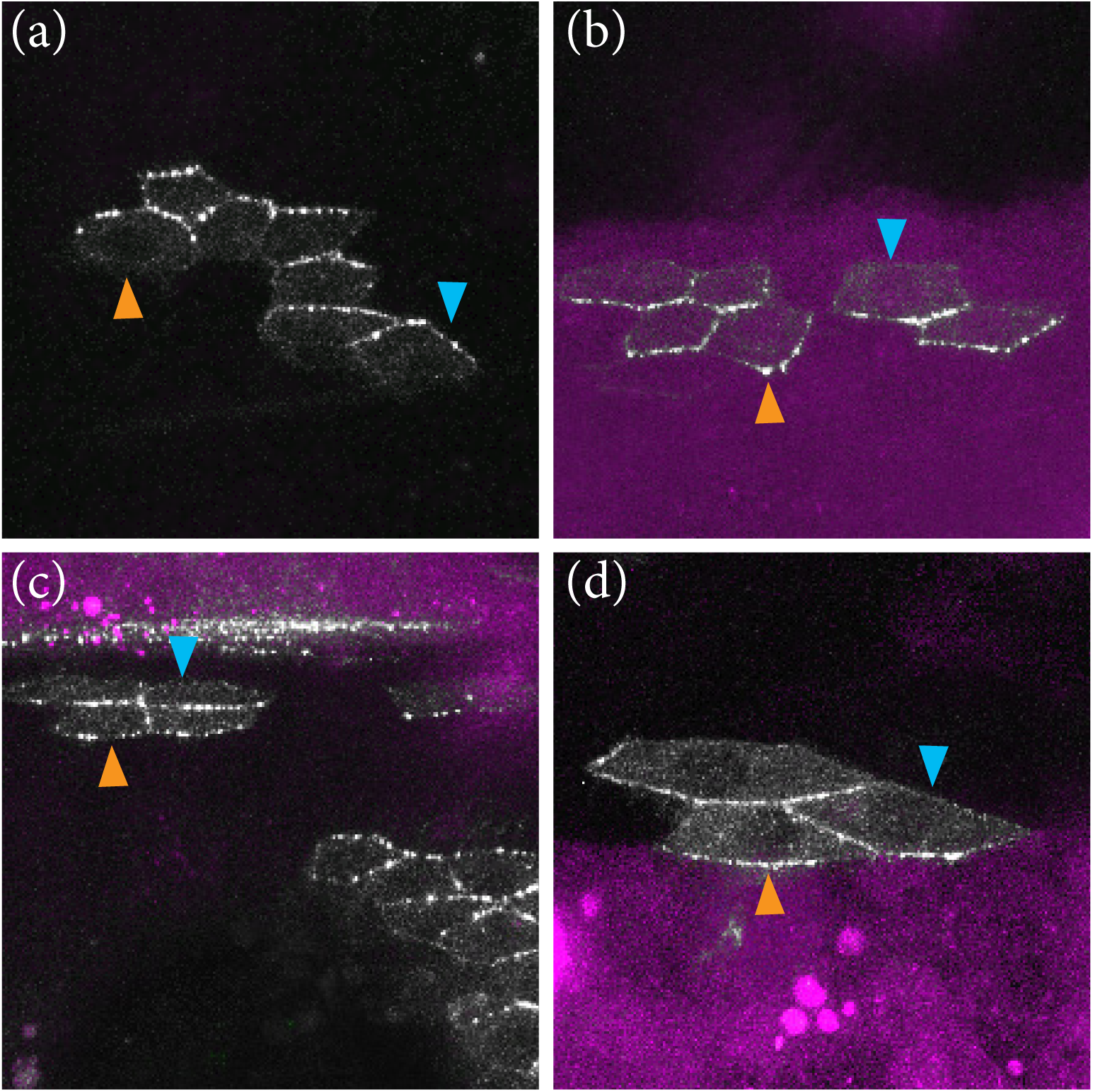
Asymmetric localisation of Dachs vis-à-vis A/P boundary. Much of the areas shown is covered with clones lacking tagged D. We see from islands of cells containing tagged D that D is located anterior in most cells of the A compartment (a) and located posterior in cells of the P compartment (b). Note that just within the A compartment, both near the anterior (c) and posterior limits (d), 1 or 2 rows of cells evince a polarity characteristic of P cells, with D located mainly on the posterior edges of the cells. The A/P boundary is demarcated by the anterior edge of *engrailed* expression that marks all cells of the P compartment (purple territory). Evidential membranes are marked with arrowheads, blue for anterior membranes and beige for posterior.

These findings place alternating fields of polarity out of register with the corresponding compartments and raise questions about the role of the compartment boundary in the genesis of polarity [cf **20**]. We therefore decided to make polarity-changing clones that overexpress *ft* in order to alter polarity near compartment boundaries. Normally such *ft*-expressing clones will reverse the polarity of surrounding cells depending on the compartment (hairs point away from the clone in the A compartment and towards it in the P). One might expect a *ft*-expressing clone located posteriorly in the A compartment and touching the compartment boundary to behave like a normal A clone at the front and reverse the polarity of wildtype cells in front of the clone, and it does so (the wildtype hairs anterior to the clone now pointing away from the clone; see figure 6a). One would expect that such a clone would meet P cells at its posterior edge and reverse the hairs behind the clone, and it does so (the wildtype hairs posterior to the clone now pointing towards the clone; see figure 6a). Correspondingly, one would expect that a clone located at the front of the P compartment and contacting the A/P boundary to reverse both at front and behind, but it fails to reverse hairs in front (figure 6b). The explanation for both types of clones is simple: because the line of polarity reversal lies anterior to the lineage boundary, an A clone contacts A cells in front of it and P cells behind.

**Figure 6.**
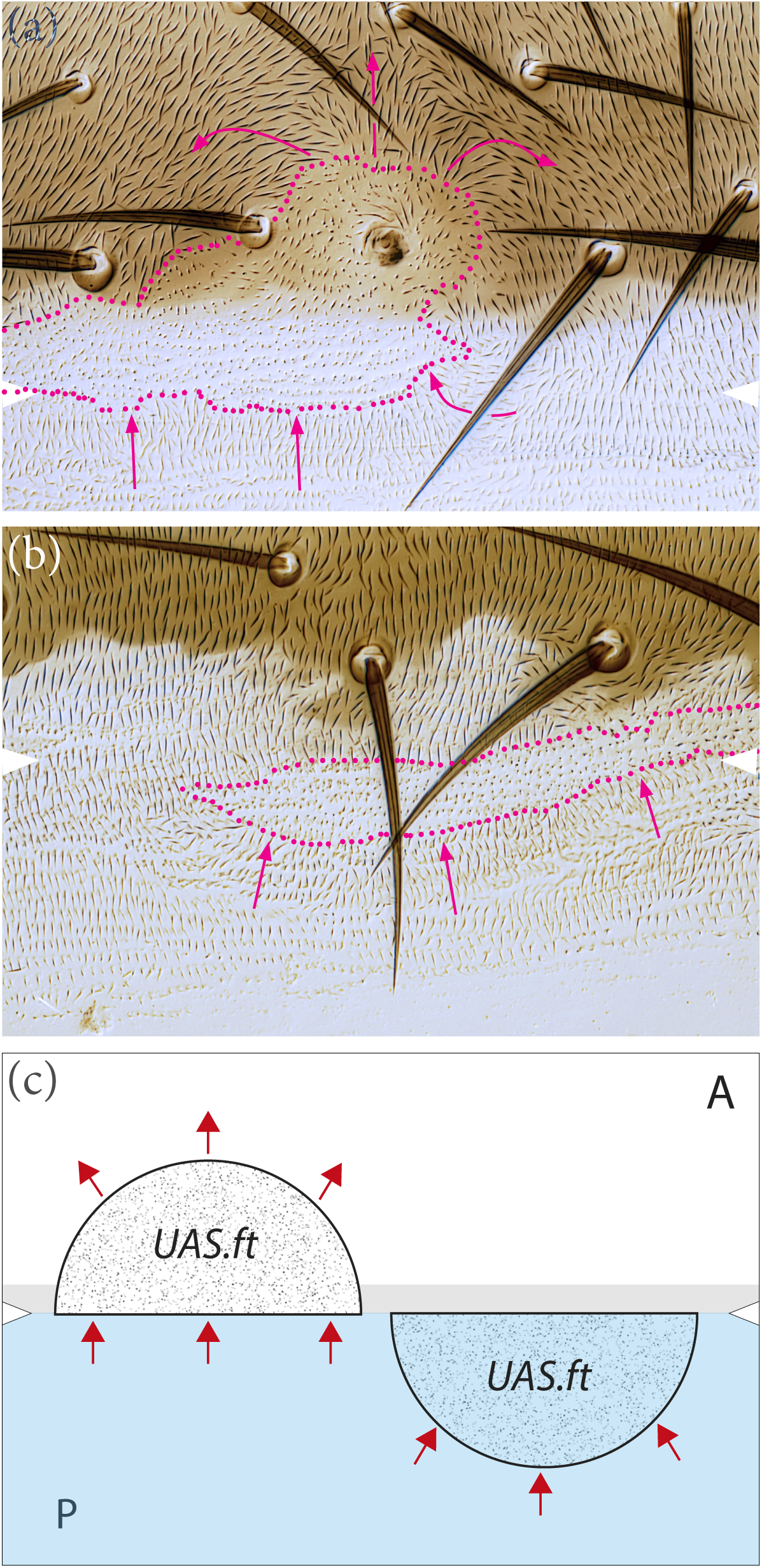
Clones overexpressing *ft* near the A/P compartment border. (a) clone of A compartment provenance, reverses territory both in front (in A territory) and behind (in P territory). (b) clone of P compartment provenance, reverses the polarity of cells behind (in P territory) but fails to reverse in front (A territory). White arrows show estimated position of A/P borders, red dots mark clone boundaries. (c) simplified diagram of above results; note line of polarity reversal shown by the location of D is anterior to the A/P boundary by 1-2 cells (shaded zone) and this explains the outcomes, see text; these results mimic those with fj- overexpressing clones [20].

However, the anterior extension of a P clone will be stopped at the A/P boundary and cannot reach the line of polarity reversal (thus it contacts cells behaving as P cells in front and therefore cannot reverse their polarity; see figure 6c).

In the flies lacking Fj, unlike in the wildtype, many cells in the anterior region of the segment show reduced polarity, with D being distributed evenly around the cell membrane. We think this finding correlates with our observation (above) that the most anterior region of the segment is where the robustness of the Ds/Ft system is most dependent on Fj. It is also pertinent that cells in the anterior region of the segment show loss of Ds asymmetry when Fj is removed (figure 3b). It also fits with the slight abdominal phenotype observed in *fj^−^* flies, in which there was some loss of normal polarity but only in the anterior region of the segment [20, 21]. Thus, all these findings point to the same conclusion: in cells of the anterior region of the segment, the polarisation by the Ds/Ft system depends more on a gradient of Fj and less on a gradient of Ds — while in the middle and rear of the A compartment the opposite is the case.

## Discussion

Here we take the familiar model system of the *Drosophila* abdomen, in this case at the pupal stage, and measure the distribution of Dachsous (Ds) in the membranes of cells *in vivo*. We describe the intercellular gradients and intracellular asymmetry across a whole metamere. Ds is distributed in a gradient which is reflexed, rising in one direction in the A compartment of each metamere and falling in the P compartment (figure 2a). Although this pattern resembles that of a morphogen, our view is that Ds/Ft is not a morphogen: the primary function of a morphogen is to provide positional information to the cells in its field [reviewed in **37, 38**], while the immediate purpose of the Ds/Ft gradient is to polarise cells. Also, and unlike an archetypal morphogen, Ds does not move from cell to cell, although the numbers of Ds molecules in the membrane of one cell affect the distribution of Ft and Ds molecules in the adjacent cell [10]. There are models of how the Ds/Ft system works, how polarity information passes from cell to cell and how a gradient of Ds activity might point the arrow of polarity [10, 13, 39, 40]. Our results validate the hypothesis that the orientation of molecular gradients determines the polarity of the cells [1, 2].

The amount of Ds forms a linear gradient along the anteroposterior axis of the A compartment, rising about 30%. We measure the cellular asymmetry in the distribution of Ds across the whole metamere. In the A compartment it is uniform and higher in the anterior membrane of the cell, that facing the bottom of the gradient, that in the posterior membrane, that facing the top of the gradient [as predicted **10**]. In the P compartment both the direction of slope of the gradient and its asymmetrical distribution in the cell are opposite to that in the A compartment [10]. The difference between anterior and posterior membranes is far less to that found when a small region of the wing imaginal disk was studied [twofold **27**]. However, modelling predicted that the difference in the primordium of the wing disc could be less than twofold [39].

### The function of Fj

There is considerable evidence from enhancer traps and from functional experiments that Fj forms a gradient [16, 20, 21]. In the A compartment of each abdominal segment it is evidenced to be highest in the most anterior region [20]. Fj phosphorylates both Ds and Ft proteins [41], the effect on Ds is to decrease its affinity for Ft, while phosphorylated Ft has an increased affinity for Ds [17, 18]. It is thought that the two opposing gradients, Ds and Fj, work together to produce asymmetric distributions of Ds-Ft bridges [10, 18]. Most pertinently, Hale et al [39] have used FRAP to investigate the stability of Ds-Ft heterodimers in the wing disk and observed differences between *fj^+^* and *fj^−^* flies. They concluded that “the overall result of removing Fj was a reduction in stability of the Ft-Ds dimer”. We cannot divine from this how the stability of bridges might impact on cell asymmetry in different parts of the abdominal segment. This is partly because Hale et al look at the conjoined membranes of two cells while we distinguish anterior from posterior membranes of each cell. We do find that removing Fj increases the relative amount of Ds in the posterior membrane over much of the A compartment. Since Ds is stable in the membrane only when joined to Ft in the next cell [14] it follows there should be more bridges in *fj^−^* segments, and, if so, how can these bridges be less stable? Hale et al [39] also deduced that the action of Fj on Ft dominated over its effect on Ds; however this finding applied to their sample area (the wing pouch) which is near the top of the Fj gradient; they do not tell us what, if any, might be the function of Fj in areas were Ds expression is high but Fj low. Our data concern the whole field and argue that Fj is essential for cellular asymmetry in only the anterior part of the A compartment (where it peaks in the wildtype), but it is also needed in the rest of the compartment to achieve a robust cellular asymmetry.

Perhaps the most problematic fact about Fj is that removing it has little overt effect on phenotype. Nevertheless, in *fj^−^* flies, we found changes in Ds distribution and a loss of robustness —shown by a reduction of the Ds/Ft system’s ability to resist polarity changes induced by clones in the Stan/Fz system. Strikingly, in the anterior ca 20% of the A compartment, the loss of Fj tends to totally depolarise the cells, eliminating the asymmetric distribution of Ds, and reducing the asymmetric localisation of D. We conclude that the main function of Fj in the abdomen, via its action on Ds and Ft, is to strengthen the Ds/Ft system mainly at the front of the A compartment where Ft is high and Ds is low.

Another clear finding demands an explanation: when the localisations of Ds in *fj^+^ and fj^−^* flies are compared, they differ considerably, but only in the posterior membranes of the cells (figure 3). It seems that Fj promotes the presence of Ds-Ft dimers more strongly in the posterior than in the anterior membranes. We offer a speculative model to explain this (figure 7).

**Figure 7.**
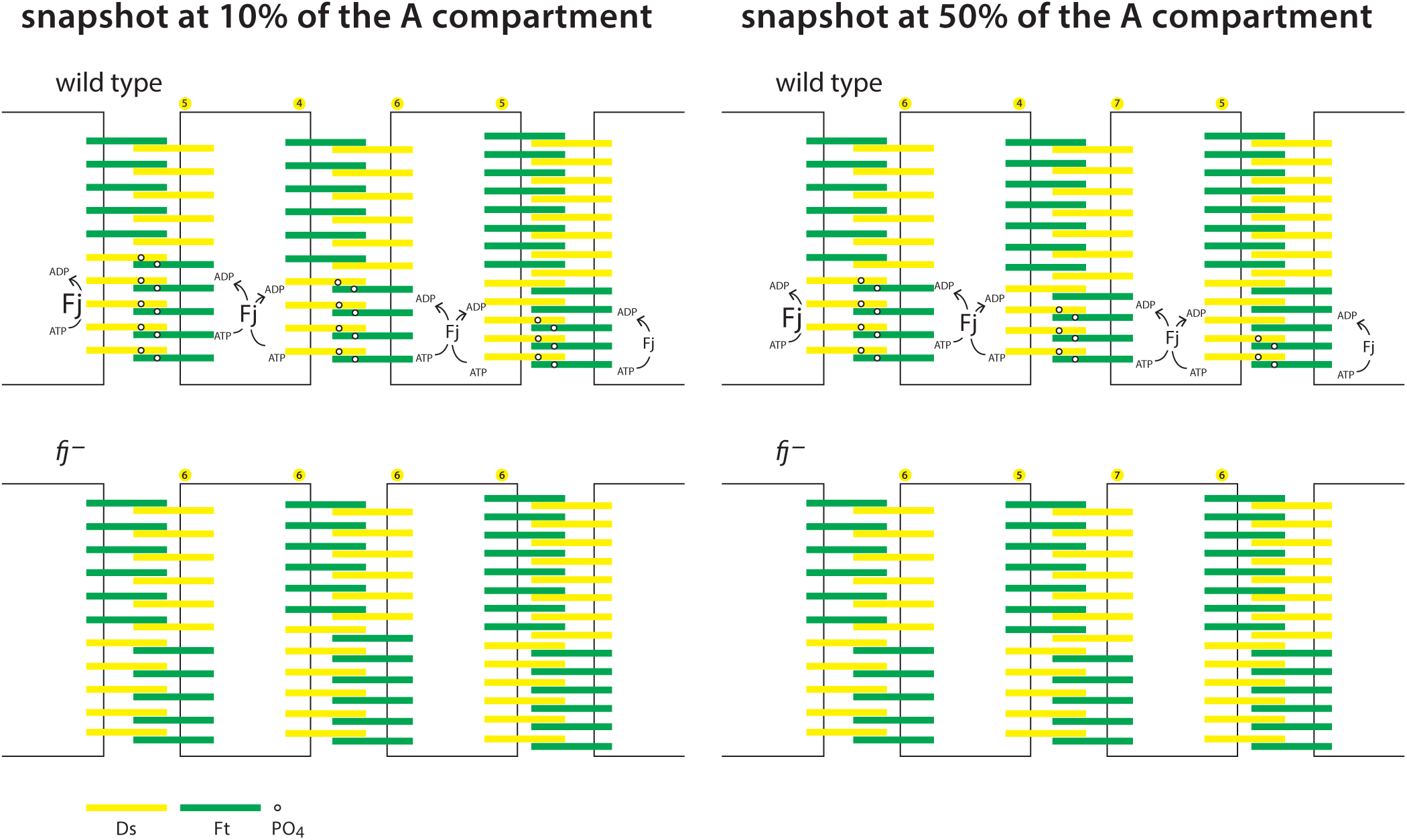
Speculative model of the action of Fj on positioning of Ds-Ft heterodimers. Why does the loss of Fj particularly affect the posterior membranes of cells? Two places in the A compartment are shown (a) 10% back from the anterior limit of the A compartment. In this area Fj is largely responsible for the supracellular gradient and cellular asymmetry. Phosphorylating Ds reduces its tendency to bind to Ft while phosphorylating Ft increases its tendency to bind to Ds [17, 18]. The Ds phosphorylated by Fj is shown to be inserted preferentially into the posterior membrane where it binds to Ft in the abutting cell. We don’t know why this might be so, it is possible that phosphorylated Ds might be transported posteriorly in the cell. Below, the same location but in *fj^−^* pupae. There is no variation in cellular asymmetry with position and the gradient is almost flat due to low levels of available Ds in this area where there is now no Fj to drive polarity. (b) 50% back from the anterior limit of the A compartment. In this area the gradient of Ds is sufficient to drive both the supracellular gradient and the cellular asymmetry, thus, below, in the absence of Fj, both the gradient and the asymmetry persist with only some reduction in strength. As with (a) we imagine phosphorylated Ds being added preferentially to the posterior membranes of cells.

### The ranges of the Ds gradients

In order to map the polarity of all the cells individually we looked at tagged D, whose asymmetric distribution depends on the localisation of Ds and Ft in the cell [32, 33]. We expected [20] and it was even reported by others [23, 35] that this inflection of cell polarity as well as the limits of the Ds/Ft gradients would coincide at the A/P borders. However, our maps of D asymmetry make clear that the changeover of polarity occurs not at the compartment border but just within the A compartment —a result that at last makes sense of earlier findings with clones overexpressing *fj*. Generally in the A compartment, hairs pointed away from the clones (suggesting a Ds gradient that rose from anterior to posterior) and, in the P compartment, hairs pointed towards the clones (suggesting a Ds gradient that rose from posterior to anterior) [20]. However, the behaviour of some *fj*-expressing clones, those that contacted a compartment border, did not fit with our expectation at that time. For example, clones belonging to the P compartment that reached the very front edge of the P compartment should reverse the polarity of A cells in front but did not do so. Why not? We were flummoxed and offered an *ad hoc* explanation [20]. Now we know that cells at the extreme rear of the A compartment have the Ds/Ft polarity of P cells, a simpler explanation makes more sense. Because *ft* and *fj*-expressing clones in the P compartment are not able to extend across the compartment boundary to contact cells with normal A polarity, they must, as observed, continue to behave as a P clones at their anterior margins because, although they confront cells of A lineage, those cells have the polarity of P cells. Consequently, effects on the disposition of bridges will reinforce, rather than alter, normal polarity.

This new picture recalls other effects of compartmental borders in fly development. An example is the A/P wing border. The interface between a signalling P and a receiving A compartment leads to a signal (Hedgehog) crossing over from P to A and initiating a response in the first cells of the receiving compartment (turning on Dpp expression, [42, 43]. We wonder if our finding relates to this: could a signal coming from the P compartment during early development initiate changes in the first cell row or two of the A compartment that spread forwards and backwards from there to induce a reflexed Ds/Ft gradient in the A and the P compartment? It is relevant that [44] have argued that the Ds/Ft gradient might be initially aligned in the anteroposterior axis in the pupal wing (that is, orthogonal to hair orientation). In which case a Ds/Ft gradient in the wing could also relate to a signal, such as Hedgehog, crossing over the boundary from the P compartment into A.

### Steepness model and growth

In thinking about the control of growth, and many have done so, we should remember that it is likely to be complicated and multifactorial. For example, regarding the role of the Ds/Ft system, removal of either Ft or Ds breaks that system and yet the flies still grow. It follows that models relating to the Ds/Ft system such as the steepness hypothesis [45, 46], or the feedforward model of growth [47] may prove insufficient. Here we find that, within each compartment, the difference between anterior and posterior membranes is largely uniform and the supracellular gradient is largely linear. These are both prerequisites for the simplest steepness model. In order to draw the arrow of PCP, anterior and posterior membranes of a cell must be locally compared and for this there is indirect evidence [48]. We conjectured that, in addition, the degree of difference between these two cell membranes might feed into the decision as to whether a cell divides or dies [45] and help to limit growth. This would amount to a dimension-sensing mechanism. The steepness model is also supported by experiments showing that interfaces between cells with different amounts of Ds lead to Hippo target genes being activated and increased local growth across that interface [49].

### One or two systems?

We have argued that the Ds/Ft and Stan/Fz systems act independently [10, 26] but this is not accepted by everyone [50]. Some authors have taken refuge in the postulate that PCP might operate differently in various organs, so the two systems might be independent in one organ (the abdomen) but are united in a single pathway of function in other organs (eg, the wing) or that any direct action of Ds/Ft on cell polarisation might constitute a “bypass pathway”). [22, 50]. We view that refuge as intrinsically precarious. The many experiments and contrasting interpretations in this area are well presented by Strutt and Strutt [51]. The recent intervention of the Pk gene into this melée has not simplified that debate [35, 52–55]. However, our results above (those comparing *fz*- expressing clones in abdomens with and without Fj, figure 4, where we ask one PCP system to act against the other), are simply explained if the systems act independently. Under the alternative model, in which the Stan/Fz system is presumed to act downstream of the Ds/Ft system, explaining the differing effects of *fz*-expressing clones on polarity in *fj^+^* and *fj^−^* flies would tax the finest minds.

Our opinion is that the two systems can function distinctly everywhere and act in conflict or in synergy. They interact late in the cellular process such as when hairs are being formed in their final orientations.

### Some remaining questions

There are many outstanding questions about the Ds/Ft system. Does the amount of Ft vary over the field? What other factors, apart from Fj, modulate the interaction of Ds and Ft molecules? How exactly is the supracellular gradient read in order to orient polarity of cells? How are opposing membranes of a cell compared in order to polarise that cell? Ds and Ft proteins are together localised into puncta [56] but why are they and are puncta required for proper function? We found the Ds/Ft gradient to be reflexed; consequently, since all the hairs point posteriorly, they must be pointing up the Ds gradient in nearly all of the A compartment and down the Ds gradient in the P compartment. How is this achieved? One simple hypothesis is that hair polarity is the outcome, in the A compartment, of both the Ds/Ft and the Stan/Fz systems instructing the hairs to point posteriorly. However, in the P compartment, the Ds/Ft system aims to point the hairs forward and the Stan/Fz aims to point backwards and, to put this too simply, the Stan/Fz system wins. The *prickle* gene also plays a part in this, see elsewhere [35, 51–54, 57].

## Data Accessibility

Data used in figures … and https://royalsocietypublishing.org/doi/10.1098/rsob.200290 - RSOB200290F8…, electronic supplementary material, figures S… can be obtained from the University of Cambridge Open Access repository (https://doi.org/10.17863/…).

## Authors’ Contributions

The experiments were conceived of by JC and PAL, all the methods devised by JC. Execution of the experiments including the making of genetic stocks depended on all the authors. The paper was written by JC and PAL. All the authors gave final approval for publication and agreed to be held accountable for their contributions.

## Acknowledgements

We thank David Strutt for his help and advice, Malcolm Burrows and Gary Struhl for encouragement and the Wellcome Trust (4 grants), the Zoology Department and the Newton Trust for supporting our experiments over the last 16 years.

## Supplementary Figures

**Figure S1.**
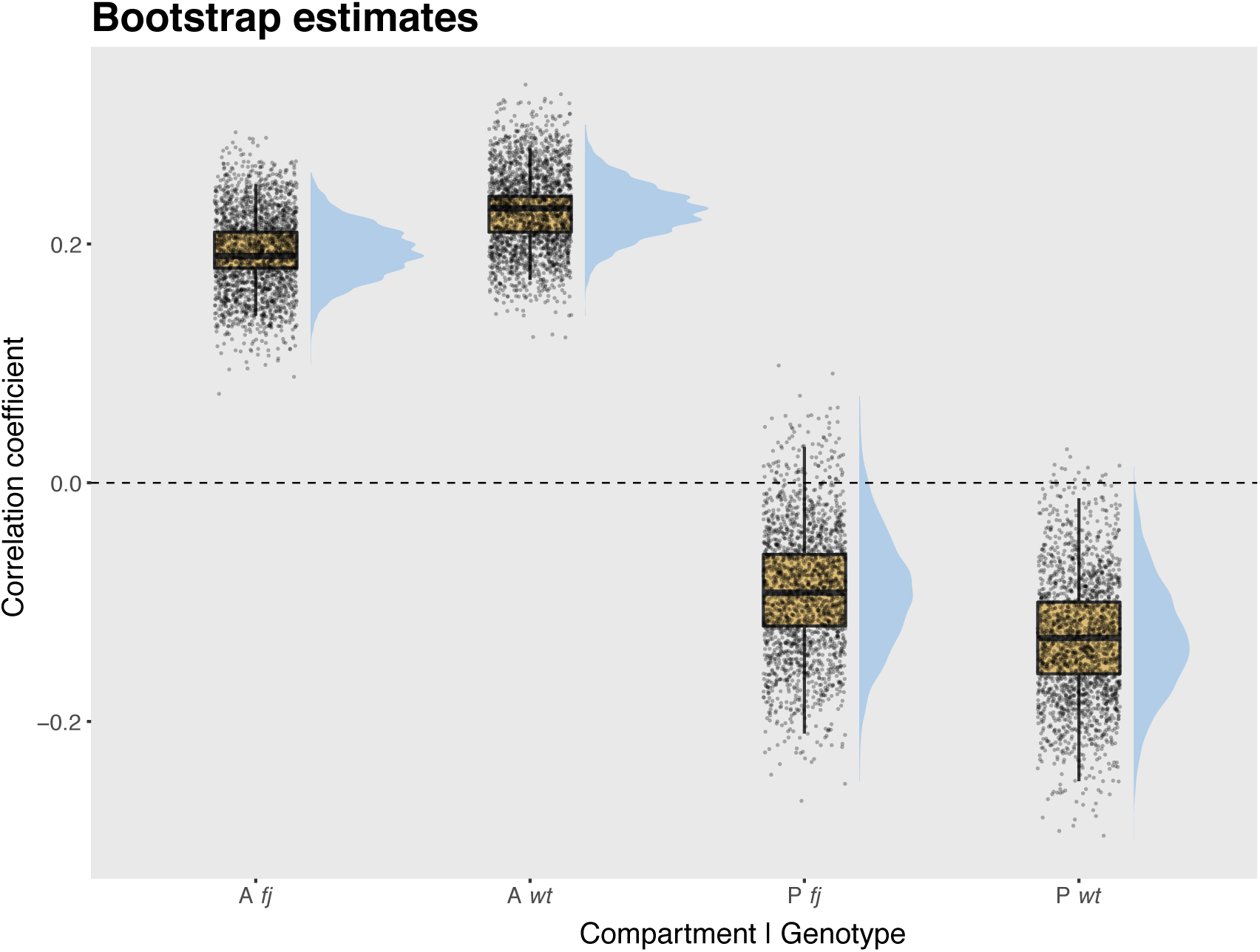
Bootstrapped estimates of the correlation between Ds accumulation at cell membranes and their position within the segment, in wiltype and *fj^−^* abdomens. Raincloud plots [58] of bootstrap estimates of correlation coefficients of the data showed in figures 2a and 3a. The estimates are clearly departing from zero in both A compartments; most of the estimates of the estimates obtained for the wildtype P compartment are negative, however, the correlation in *fj^−^* compartments appear to be lost.

**Figure S2.**
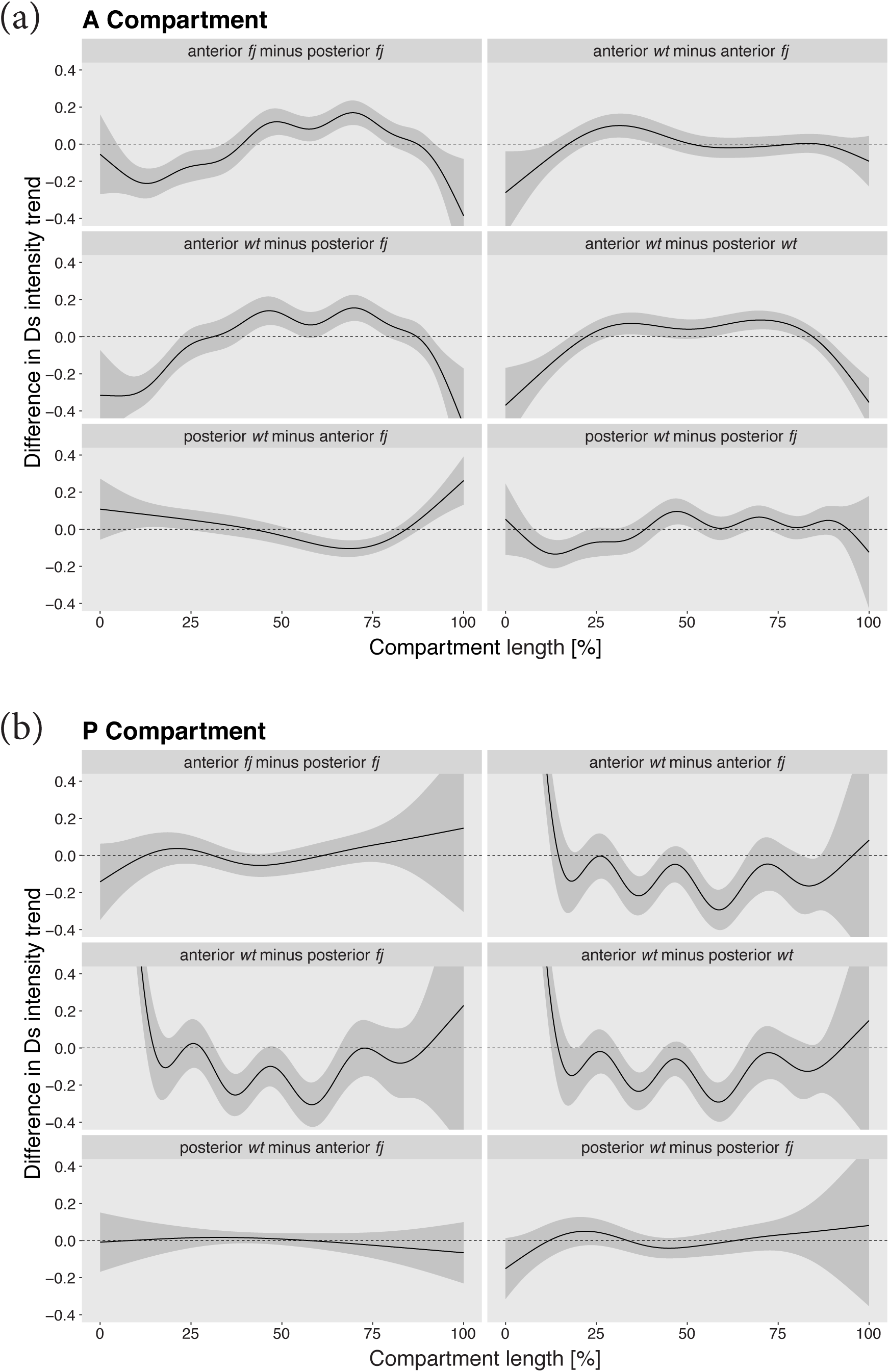
Estimated differences of trends in Ds accumulation. Differences of trends in Ds accumulation of pairs of smooths in anterior and posterior cell membranes of the A compartments (a), and anterior and posterior cell membranes of the P compartments (b) of wild type and *fj^−^*.

**Figure S3.**
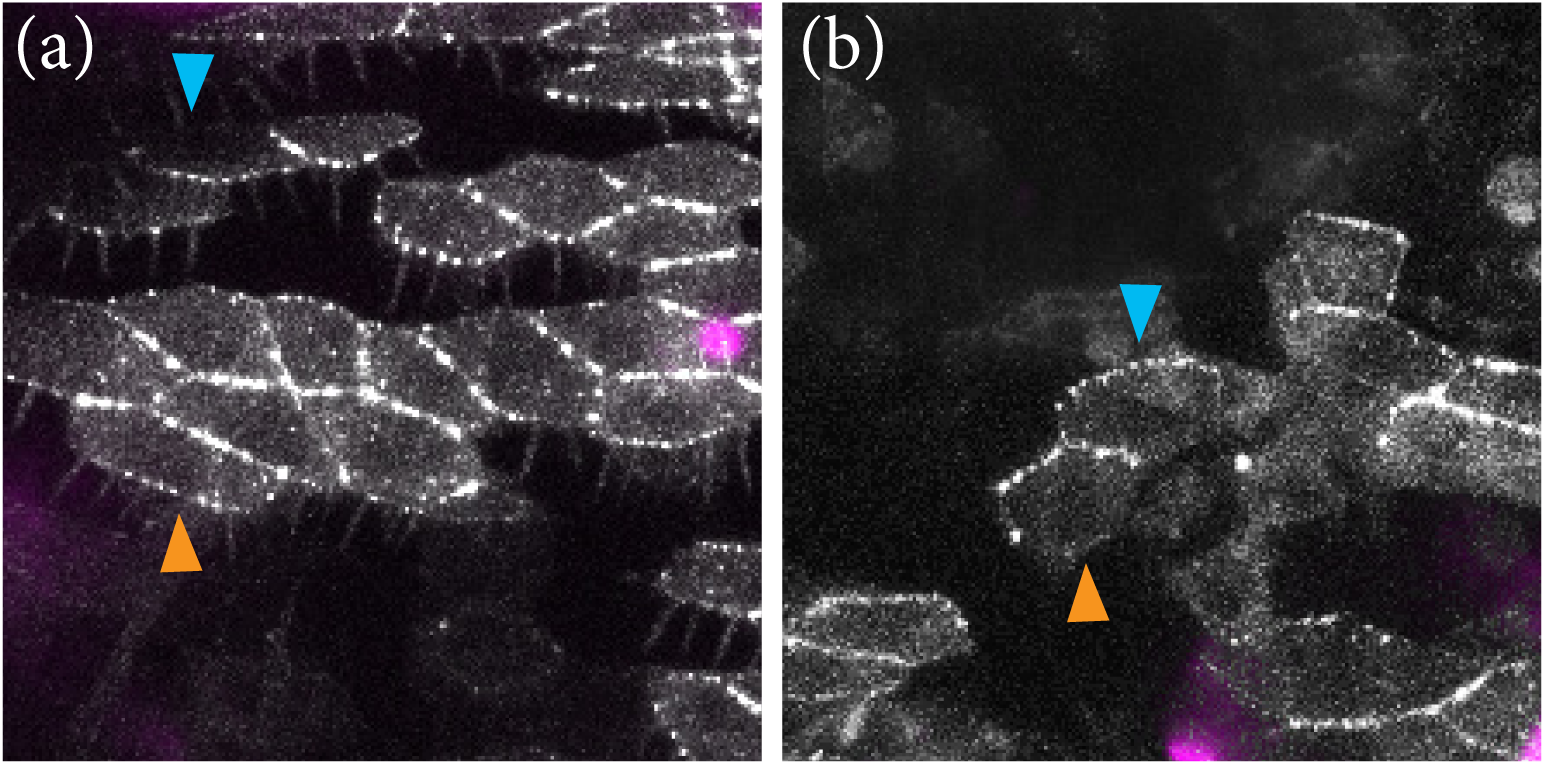
The location of D in *fj^−^* pupae. (a) Images shows an anterior region of A compartment; in the wildtype all cells of this region show D localised anteriorly. But in fj- cells show either posterior localisation of D (the anteriormost cells in the image) or variable or unclear asymmetry. (b) An area near the middle of the A compartment showing that D, as in wildtype cells in this region, is located anteriorly. Blue arrow marks an anterior membrane, with no D visible and orange arrow marks the posterior membrane of a cell with some D posteriorly.

**Figure S4.**
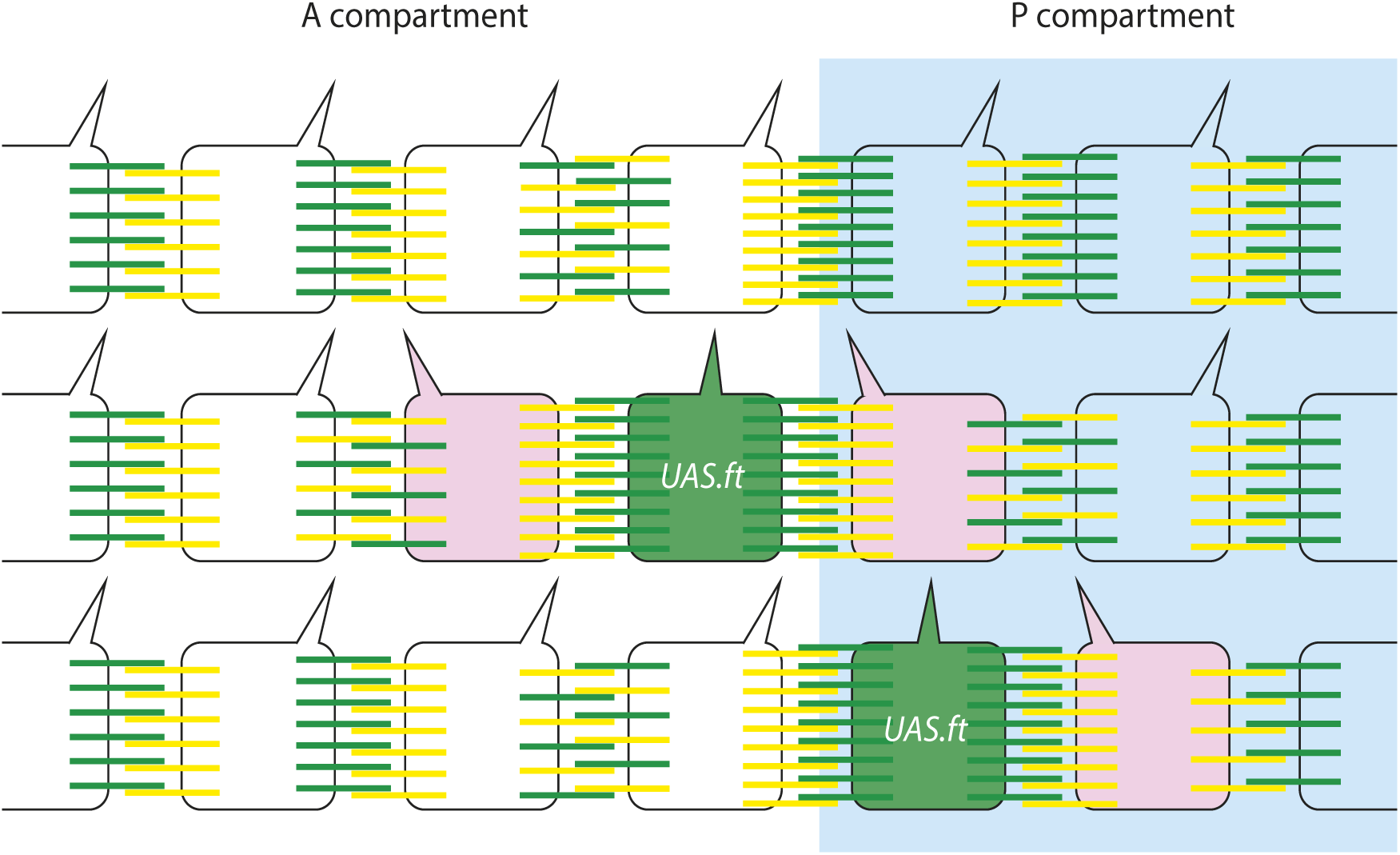
T**o illustrate effects of *ft* overexpressing clones on bridges and polarity** Upper row shows wildtype. Normally, in the A compartment most Ds is on the anterior membrane (see figure 2) but the rearmost cell of the A compartment has reversed Ds/Ft polarity (as shown by the localisation of D, see figure 5), with most Ds on its posterior membrane. In the P compartment there is more Ds on the posterior membrane Middle row shows a clone overexpressing ft at the rear of the A compartment. Cells of this clone (one cell is shown) draw Ds to the adjacent membranes on both neighbours and both these neighbours polarities are reversed (cells labelled in pink). Bottom row shows a clone overexpressing ft at the front of the P compartment. Cells of this clone draw Ds to the adjacent membranes on both neighbours but only its neighbour in the P compartment has its polarity reversed (cells labelled in pink). The neighbour in the A compartment retains its normal normal polarity (see Figure 6).

